# An enduring role for hippocampal pattern completion in addition to an emergent non-hippocampal contribution to holistic episodic retrieval after a 24-hour delay

**DOI:** 10.1101/2023.09.15.557911

**Authors:** Bárður H. Joensen, Jennifer E. Ashton, Sam C. Berens, M. Gareth Gaskell, Aidan J. Horner

## Abstract

Episodic memory retrieval is associated with the holistic neocortical reinstatement of all event information; an effect driven by hippocampal pattern completion. However, whether holistic reinstatement occurs, and whether hippocampal pattern completion continues to drive reinstatement, after a period of consolidation is unclear. Theories of systems consolidation predict either a time-variant or -invariant role of the hippocampus in the holistic retrieval of episodic events. Here, we assessed whether episodic events continue to be reinstated holistically and whether hippocampal pattern completion continues to facilitate holistic reinstatement following a period of consolidation. Female and male human participants learnt ‘events’ that were composed of multiple overlapping pairs of event elements (e.g., person-location, object-location, location-person). Importantly, encoding occurred either immediately before or 24-hours before retrieval. Using fMRI during the retrieval of events, we show evidence for holistic reinstatement, as well as a correlation between reinstatement and hippocampal pattern completion, regardless of whether retrieval occurred immediately or 24-hours after encoding. Thus, hippocampal pattern completion continues to contribute to holistic reinstatement after a delay. However, our results also revealed that some holistic reinstatement can occur without evidence for a corresponding signature of hippocampal pattern completion after a delay (but not immediately after encoding). We therefore show that hippocampal pattern completion, in addition to a non-hippocampal process, has a role in holistic reinstatement following a period of consolidation. Our results point to a consolidation process where the hippocampus and neocortex may work in an additive, rather than compensatory, manner to support episodic memory retrieval.

## Introduction

The recollection of episodic events is holistic in nature (Yonelinas, 1994; Tulving, 1983); underpinned by a hippocampal pattern completion process (Hopfield, 1982; Marr, 1971; McClelland et al., 1995; Norman & O’Reilly, 2003; Treves & Rolls, 1992) that drives the reinstatement of event information in the neocortex (Johnson et al., 20009; Wheeler et al., 2000; Woodruff et al., 2005).

Evidence for hippocampal pattern completion and holistic neocortical reinstatement has been shown immediately following encoding (Grande et al., 2019; Horner et al., 2015). However, we do not yet know whether this relationship changes following a delay. This is a critical question as standard theories of consolidation predict a time-limited role for the hippocampus (McClelland et al., 1995; Squire & Alvarez, 1995), while others argue that the hippocampus is always required for episodic retrieval (Nadel & Moscovitch, 1997; Sekeres et al., 2018; Winocur & Moscovitch, 2011).

There is evidence for decreases in (Niki & Lao, 2002; Takashima et al., 2006; Takashima et al., 2009) and sustained (Addis et al., 2004; Bonnici et al., 2012; Gilboa et al., 2004; Maguire et al., 2001; Ryan et al., 2001) hippocampal activity during retrieval following a delay (see also Ritchey et al., 2015). Critically, most previous studies have compared hippocampal activity (or multivariate patterns) at immediate and delayed retrieval, and therefore have not directly assessed whether hippocampal pattern completion continues to mediate the reinstatement of episodic events in the neocortex. Importantly, changes in univariate activity over time could be driven by different mechanisms not related to systems consolidation, such as differential rates of forgetting for independent neocortical and hippocampal representations formed at encoding (Andermane et al., 2021; Fisher & Radvansky, 2018; Sekeres et al., 2016). Thus, by directly assessing the relationship between hippocampal pattern completion and neocortical reinstatement across delay, we can more directly assess the mechanistic predictions of systems consolidation theories.

Here we used an experimental assay of episodic memory (Horner & Burgess, 2014; Horner et al., 2015) that allowed us to assess the delay-(in)variant role of hippocampal pattern completion in neocortical reinstatement. Across two encoding sessions, separated by ∼24-hours, participants learnt three-element events. Events were learnt over two or three separate encoding trials, with each trial presenting one of three possible pairs for an event. Events had different associative structures: ‘closed-loops’ where all three pairs were encoded and ‘open-loops’ where only two of the three pairs were encoded. At retrieval, we tested each of the learnt pairs from each event, with half of the events learnt ∼24 hours before retrieval and the other half learnt immediately before retrieval.

We identified neocortical regions that showed differences in BOLD activity between elements that were the cue or target of retrieval and those that were non-targets. Greater activity for closed-vs open-loops in non-target regions was then used as a marker of holistic neocortical reinstatement (Horner et al. 2015). Next, we correlated non-target neocortical reinstatement with a signature of hippocampal pattern completion (closed-vs open-loop hippocampal activity), for events encoded immediately and ∼24 hours earlier. We used a general linear mixed-effect model to assess both the slope and the intercept of the relationship between the hippocampal contrast and neocortical reinstatement.

A slope greater than zero would indicate a positive correlation, consistent with hippocampal pattern completion contributing to holistic neocortical reinstatement. If the steepness of the slope is consistent across delay, this would indicate a continued role for hippocampal pattern completion, while a shallower (or zero) slope in the delay relative to immediate condition would suggest a time-limited role. Critically, in the presence of a positive slope, a zero intercept would suggest that holistic reinstatement only occurs in the presence of hippocampal pattern completion. An intercept greater than zero would indicate that some holistic reinstatement can occur without a corresponding signature of pattern completion. This would point to an emergent non-hippocampal contribution that is additive (rather than compensatory) to the role of hippocampal pattern completion in holistic reinstatement.

## Methods

### Participants

An effect size *d* = .67 for detecting a significant non-target reinstatement effect was calculated from previous work (Horner et al., 2015) with *n* = 26. Cohen’s *d* was calculated from the reported *t*-statistic divided by the square root of the within-subject sample size (Lenth, 2006). As forgetting is expected to occur when retrieval follows a ∼24 hours delay, retrieval accuracy is expected to be lower than in Horner et al. (2015). Pilot data collected with *n* = 12, revealed ∼14% decrease in retrieval accuracy following a 24-hour delay, leading to an adjusted effect size *d* = .62. Using G*Power (Faul et al., 2009) and the adjusted effect size *d* = .62, we estimated that a sample size of *n* = 30 would be required to detect a significant non-target reinstatement effect, if one is present, at a power of .90 and *α* = .05.

Given that previously reported effect sizes are likely to be overestimates of the true effects size, and because we wanted to compare neocortical reinstatement as well as hippocampal-neocortical correlations between two conditions, we aimed to obtain 50 useable datasets (upper limit set by resource constraints). An initial sample of *n* = 30 was collected and analysed (due to time constraints in relation to submitting a PhD thesis; reported in Joensen, 2019) before the final sample was collected.

57 right-handed participants were recruited from the University of York student population. All participants gave written informed consent and took part in exchange for course credit or monetary compensation. Participants had either normal or correct-to-normal vision and reported no history of neurological or psychiatric illness. Data from seven participants could not be included in the final sample due to motion related artefacts in the imaging data (*n* = 1), scanner faults resulting in no acquired imaging data (*n* = 2) or below chance memory performance (accuracy < ∼16.7%) (*n* = 4). Accordingly, the analyses included 50 participants (39 female, 11 male) with a mean age of 21.6 years ± 4.43 *SD*. The experiment was approved by the York Neuroimaging Centre (YNiC) ethics committee, University of York, UK.

### Materials

The stimuli consisted of 72 locations (e.g., *kitchen*), 72 famous people (e.g., *Barack Obama*), and 72 common objects (e.g., *hammer*). From these, 72 random location-person-object events were generated for each participant. Note that we used the term ‘event’ to refer to the three elements (location, person, object) that were assigned to the same associative structure (closed-or open-loop), leaving aside any considerations as to how these relate to real-world events (but see Horner et al., 2015 for discussion on this point).

Each of the 72 events were assigned to the within-subject conditions of Loop (open-loop or closed-loop) and Delay (delay or no-delay). Thus, 18 different events were randomly assigned to either: (1) open-loop, delay; (2) closed-loop, delay; (3) open-loop, no-delay; or (4) closed-loop, no-delay. For closed-loops, all three possible pairwise associations for each event were encoded, while for open-loops, only two out of the three possible pairwise associations were learnt. Events were never presented together at encoding or retrieval; only specific pairwise associations were encoded and retrieved for each event, depending on whether they were open-or closed-loop. Note that open-loops are equated to closed-loops in the number of elements, but not in the number of pairs. We have previously shown (Horner et al., 2015; Horner & Burgess, 2014) that dependency is not seen for open-loops when three overlapping pairs are encoded as an associated chain (e.g., *kitchen-hammer, kitchen - Barack Obama, Barack Obama - dog*), controlling for the number of associations (but not the number of elements). Any differences in retrieval between the two conditions in the current experiment are therefore unlikely to be driven by differences in the number of associations.

### Procedure

The experiment consisted of two encoding sessions and a single retrieval session. Both encoding sessions took place outside the scanner and were separated by ∼24 hours (*M ± SD* = 23 hours, 54 minutes ± 130 minutes), including overnight sleep. The retrieval session took place inside the scanner immediately after the second encoding session.

#### Encoding

During encoding, participants were presented with specific pairwise associations from each of the 72 events. Participants learnt one pairwise association per encoding trial. All pairwise associations were presented on a computer screen as words, with one item to the left and one item to the right of fixation (see Figure 1A). The left/right assignment was randomly chosen on each trial. The words remained on screen for 6s. Participants were instructed to imagine, as vividly as possible, the elements interacting in a meaningful way for the full 6s. Each word-pair presentation was preceded by a 500-ms fixation and followed by a 500-ms blank screen. For 36 out of the 72 events, participants encoded all three possible pairwise associations; forming closed-loops (see Figure 1C). For the remaining 36 triplets, participants encoded two out of the three possible pairwise associations; forming open-loops (see Figure 1D). 18 closed-and 18 open-loops were randomly assigned to each of the encoding sessions, resulting in a total of 90 encoding trials per encoding session (i.e., 54 closed-loop and 36 open-loop trials per session). Participants were not told that any pairwise associations would overlap with each other at encoding nor were they told about the open- and closed-loop structures.

**Figure 1.**
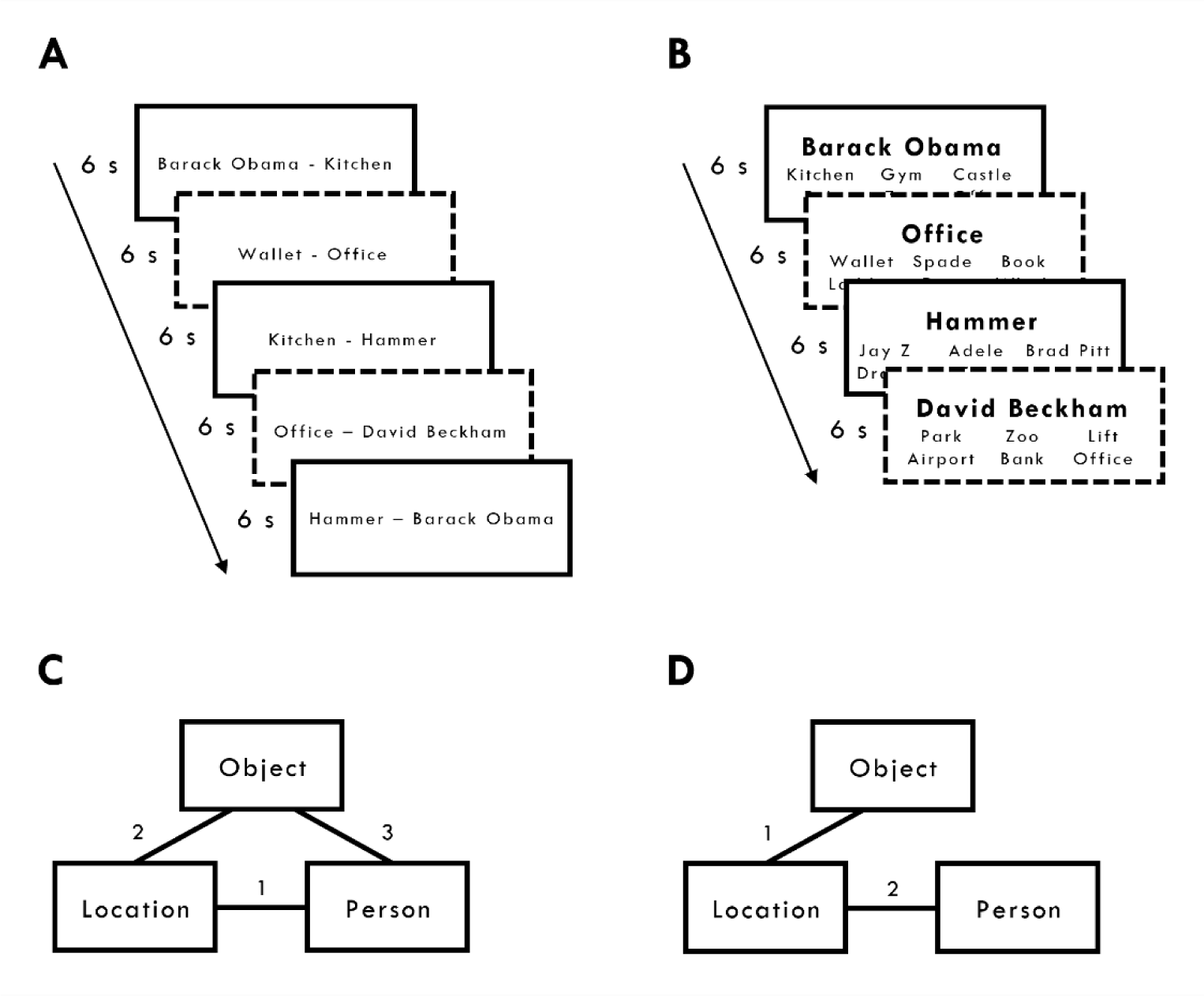
Experimental design. **(A)** Encoding. Participants saw multiple pairwise associations. They imagined each association ‘interacting in a meaningful way as vividly as possible’ for 6s. Each association was preceded by a 500-ms fixation and followed by a 500-ms blank screen. Participants encoded open- and closed-loop pairwise associations in an intermixed manner. In the open-loop condition, participants did not encode the third and final association (e.g., wallet-Beckham) (see D). Solid and dotted lines were not presented but highlight closed-(solid lines) and open-loops (dotted lines). **(B)** Retrieval. Participants were presented with a single cue and required to retrieve one of the other elements from the same event from among five foils (elements of the same type from other events) in 6s. Each cued-recognition trial was preceded by a 1s fixation and followed by a 1s blank screen. **(C)** The associative structures of closed-loops with example encoding order for three pairwise associations (numbers 1-3). **(D)** The associative structure of open-loops with example encoding order for the two pairwise associations (numbers 1-2). The third and final association (i.e., person-object in this example) was not shown to the participants.

Each encoding session consisted of three blocks of 18, 36 and 36 trials, respectively, with one pairwise association from each event presented during each block. Each encoding session contained novel triplets for the open- and closed-loop conditions that were not presented in the other encoding session. During the first encoding block, only pairwise associations for closed-loops were presented. This ensured that the duration between encoding the last pairwise association and retrieval was consistent for closed- and open--loops from each encoding session. In blocks two and three, the open- and closed-loop associations were randomly presented in an intermixed manner. The presentation order was randomised for each block. A break of 10s followed every 18 encoding trials.

Each open-loop consisted of a common element (e.g., if participants encoded location--person, and location-object pairs, then location would be the common element). For each encoding session, six out of the total 18 open-loops were pseudo-randomly assigned to one of the three possible common elements (i.e., location, people or objects). The presentation order for the open-loops across encoding blocks two and three was: (1) person-location, location-object; (2) location-object, object-person; (3) object-person, person-location. The presentation order for the closed-loops across the three encoding blocks was: (1) person-location, location-object, object-person; (2) location-object, object-person, person-location; (3) object-person, person-location, location-object. These counterbalanced orders ensured that the first two encoding trials for open-loops and closed-loops were identical in structure, such that closed-loops only differed in relation to the presentation of a third encoding trial.

#### Retrieval

During retrieval, participants performed a six-alternative forced-choice cued-recognition task. On each trial, a cue and six potential targets were presented simultaneously on screen. The cue was presented in the middle of the screen with the six possible targets; one target and five foils form the same category (e.g., if the target was *hammer*, the five foils would be other randomly selected objects from the other events, regardless of closed- vs open-loop or delay vs no-delay status) presented in two rows of three below the cue (see Figure 1B). Participants had a maximum of 6s to respond with a button response that corresponded to the targets position on screen and were instructed to be as accurate as possible in the time given. The position of the correct target was randomly selected on each retrieval trial. The cue and six targets were presented until a response was made or when the maximum 6s limit was reached (*M* response times *± SD* = 2.86 s ± 0.42 s). Missing responses (i.e., responses that fell outside the 6s response window) were treated as incorrect trials for both accuracy and dependency (*M* percentage of missing responses *± SD* = 3.54% ± 3.22%). A 2×2 (Loop x Delay) ANOVA, where the dependent variable was the proportion of nonresponses, showed no significant effects (*F*s < .37). Thus any differences in dependency across conditions are unlikely to be caused by assuming missing responses would have been incorrect. Note also that due to the 6-alternative forced choice recognition test, the chance of guessing correctly was relatively low (∼16.7%).

Each encoded pairwise association (regardless of Delay or Loop status) was tested in both directions (e.g., cue: location, target: person; cue: person, target: location) across two scanning runs (total retrieval trials = 360; 180 trials per run). For the open-loops we only tested the pairwise associations that were directly encoded - and not those that could potentially be inferred from overlapping nature of pairwise associations (e.g., Shohamy & Wagner, 2008; Zeithamova et al., 2012) - resulting in four retrieval trials per open-loop compared to six retrieval trials per closed-loop. The presentation order was optimised to measure univariate BOLD activity in each of the four within-subject experimental conditions (i.e., open-loop delay, open-loop no-delay, closed-loop delay, closed-loop no-delay) (optimisation algorithm available at https://osf.io/eh78w/). To avoid adaptation effects, cue-target associations from the same closed- or open-loop were never presented on successive trials and each block contained 18 null trials that each lasted 6s. In each retrieval block (or scan run), cue-target associations belonging to nine closed- and open--loops encoded during the first encoding session, and nine closed- and nine open-loops encoded immediately before retrieval were tested in a random order, making a total of 180 retrieval trials, in addition to 18 null trials. Each trial was preceded by a 1s fixation and followed by a 1s blank screen.

### Modelling retrieval dependency

To model retrieval dependency, 2×2 contingency tables for the observed data and independent model (see below) were created for each participant to assess the proportion of joint retrieval and joint non-retrieval between the retrieval of two elements (e.g., person, object) when cued by a common element (e.g., location) (A_B_A_C_), and between the retrieval of a common element (e.g., location) when cued by the other two elements from the same (e.g., person, object) (B_A_C_A_). For the closed-loops, six contingency tables were constructed (as we test all three pairwise associations in both directions) and for the open--loops, four contingency tables were constructed (as we only tested the two directly encoded pairwise associations in both directions).

We also constructed contingency tables that estimate the number of trials that would fall into each of the four cells if the retrieval of pairwise associations within an open- or closed-loop was independent. This independent model assumes that pairwise associations from a given event are retrieved independently of one another - i.e., if a participant retrieved one pairwise association from an event (un)successfully this does not predict the participants ability to (un)successfully retrieve another pairwise association from the same event.

The 2×2 contingency tables for the data show the number of events that fall within the four cells (i.e., for the A_B_A_C_ analysis, both A_B_ and A_C_ correct, A_B_ incorrect and A_C_ correct; A_B_ correct and A_C_ incorrect; and both A_B_ and A_C_ incorrect, where A_B_ = cue with location (A) and retrieve person (B) and similarly for A_C_, where C refers to object). The table for the independent model (Table 1) shows the predicted number of events that fall in the four cells, given a participant’s overall level of accuracy, when the retrieval of within-event associations is assumed to be independent.

**Table 1.** Contingency table for the independent model for correct and incorrect retrieval, over *N* events (*i* = 1 to N), for elements B and C when cued by A.

| Retrieval of Element C | Retrieval of Element B |  |
| --- | --- | --- |
| | Correct ($P_{AB}$) | Incorrect ($1 - P_{AB}$) |
| Correct ($P_{AC}$) | $\sum_{i=1}^N = P_{AB_i} P_{AC_i}$ | $\sum_{i=1}^N = P_{AC_i} (1 - P_{AB_i})$ |
| Incorrect ($1 - P_{AC}$) | $\sum_{i=1}^N = P_{AB_i} (1 - P_{AC_i})$ | $\sum_{i=1}^N = (1 - P_{AB_i}) (1 - P_{AC_i})$ |

For a given participant, the proportion of correct retrievals of, for instance, element B when cued by A is denoted by *P*_AB_ (i.e., the mean performance for B when cued by A across all events). For the independent model, when cued by A, the probability of (1) correct retrieving B and C (across all events) is equal to *P*_AB_ *P*_AC_; (2) correctly retrieving B but not C is equal to *P*_AB_ (1 - *P*_AC_); (3) correctly retrieving C but not B is equal to *P*_AC_ (1 - *P*_AB_); and (4) incorrectly retrieving both B and C is equal to (1 - *P*_AB_) (1 - *P*_AC_).

Once the contingency tables for the data and the independent model were constructed, the proportion of joint retrieval and joint non-retrieval were calculated for each contingency table separately, by summing the leading diagonal cells and dividing by the total number of events (i.e., the proportion of events where two overlapping pairwise associations within an event are both retrieved either correctly or incorrectly). This measure was then averaged across the six or four contingency tables (dependent on whether the tables related to open- or closed-loops) to provide a single measure of the proportion of joint retrieval and non-retrieval for the data and independent model separately. For brevity, we refer to this measure as the ‘proportion of joint retrieval’, but note that it includes both the proportion of joint retrieval and joint non-retrieval.

This proportion of joint retrieval measure scales with accuracy. We therefore compare this measure in the data relative to the independent model. As such, the independent model serves as a lower bound that can be compared with the proportion of joint retrieval in the data. We therefore have a single measure of ‘retrieval dependency’ (i.e., the difference between the proportion of joint retrieval in the data relative to the independent model) for each participant and condition.

### Behavioural analyses

A 2×2 (Loop x Delay) ANOVA was used for the behavioural analysis of retrieval accuracy and retrieval dependency, with the within-subject factor Loop referring to whether the events formed closed- or open-loops, and Delay referring to whether closed- or open-loops were encoded immediately (no-delay) or 24 hours prior to retrieval (delay). For the analysis of retrieval dependency, we also report one sample *t*-tests comparing retrieval dependency in each condition with zero. Retrieval dependency significantly greater than zero provides evidence for the presence of dependency. Alpha was set to .05 (two-tailed) for all statistical tests. For each ANOVA, a *η_p_^2^* effect size is reported, and for *t*-tests we report a Cohen’s *d* effect size reflecting the mean difference between conditions divided by the standard deviation of the difference scores. All analyses were conducted using JASP (JASP Team, 2022).

### fMRI acquisition

All functional and structural volumes were acquired on a 3-Tesla Siemens MAGNETOM Prisma scanner equipped with a 64-channel phase array head coil. T2*-weighted slices were acquired with echo-planar imaging (EPI). 48 axial slices (∼0° tilt to the AC-PC lines) per volume were acquired in an interleaved order with the following parameters: acquisition matrix = 64 x 64, repetition time = 1200ms, echo time = 26ms, flip angle = 75°, slice thickness = 3mm, in-plane resolution = 3 x 3mm, multi-band acceleration factor = 2. To allow for T1 equilibrium, the first three EPI volumes were acquired prior to the task and then discarded. As the retrieval phase varied in length across participants (as each cued-recognition trial was self-paced, up to a maximum of 6s per trial), the number of acquired volumes differed across participants. Note that the two retrieval blocks corresponded to two separate functional scanning runs. The mean number of volumes acquired during retrieval was 796.80 (range 621 - 955) and 774.04 (range 616 - 942) for the first and second retrieval block, respectively. A field map was acquired to allow for the correction of geometric distortions due to field inhomogeneities. For the purpose of co-registration and image normalisation, a whole-brain T1-weighted structural scan was acquired with a 1mm^3^ resolution using a magnetisation-prepared rapid gradient echo (MPRAGE) pulse sequence.

### fMRI analyses

#### Pre-processing

Image pre-processing was performed in SPM12. EPI images were corrected for field inhomogeneity based on geometric distortions, and spatially realigned to the first image of the time series. EPI images were spatially normalised to MNI space with transformation parameters derived from warping each participant’s structural image to a T1-weighted average template image (using the DARTEL toolbox, Ashburner, 2007). EPI images were spatially smoothed with an isotropic 8mm FWHM Gaussian kernel prior to analysis.

#### General analysis

At the first level, BOLD activity was modelled by a boxcar function from cue/target onset to the time a response was made (up to a maximum of 6s). The predicted BOLD response was then convolved with a canonical hemodynamic response function to produce regressors of interest. In addition to the main regressors of interest, all first-level models included a set of nuisance regressors: six movement parameters, their first-order derivatives, and volume exclusion regressors censoring periods of excessive motion. The volume exclusion regressors were defined as volumes where the movement derivatives exceeded 1.5mm translation (i.e., 0.5 x slice thickness) or 1° rotation (*M* number of excluded volumes ± *SD* = 0.79 ± 3.60). Parameter estimates for each regressor of interest were included in the second level analysis to identify consistent effects across participants. Unless otherwise stated, all second level models explicitly modelled subject effects. All effects reported outside the hippocampus are *p_FWE_* < .05 family-wise error (FWE) whole-brain corrected (cluster size > 30 voxels). We also performed small-volume corrected (SVC) analyses in the hippocampus given the *a priori* predictions related to this brain region. This mask for SVC was created using the WFU PickAtlas toolbox (Maldjian et al., 2003) with the hippocampus defined from the Automated Anatomical Labelling atlas (Tzourio-Mazoyer et al., 2002).

#### Neocortical reinstatement

The first level model included 24 regressors that related to all cue-target pairs (i.e., person-location, person-object, location-person, location-object, objects-person and object-location) for each of the four experimental conditions (open vs closed and delay vs no-delay). At the second level, regions that showed greater BOLD response to cueing and retrieving each element-type (i.e., cue/target), relative to when an element-type was not cued or required to be retrieved (i.e., non-target), collapsed across the four experimental conditions were identified. This revealed three separate regions of interest across the three element-types (see Table 3). For each region of interest - defined as a 9mm radius sphere centred on the peak coordinate - BOLD responses for each individual across all 24 regressors were extracted. The BOLD response when the element associated with a given region was either the cue, target, or non-target across the four experimental conditions was then calculated. As such, estimates of the BOLD response across 12 conditions (i.e., cue, target, and non-target x closed- and open-loop x delay and no-delay) were obtained for each of the three regions of interest.

#### Non-target reinstatement and correlated activity

The first level model included four regressors corresponding to trials from each of the four experimental conditions (i.e., open-loop, delay; closed-loop, delay; open-loop, no-delay; and closed-loop, no-delay). At the second level, two separate random-effects models were created, one for the delay and one for the no-delay condition. Each model included a single regressor corresponding to the contrast between closed- and open-loops (i.e., open-loop vs closed-loops for the respective delay or no-delay condition). This was calculated by taking the mean difference in BOLD response between the closed- and open-loop conditions for the non-target element across the three regions of interest. Given the prior hypothesis regarding the hippocampus, statistical effects in the bilateral hippocampus are *p* < .05 FWE SVC within a bilateral hippocampal mask.

## Results

### Behaviour

#### Retrieval accuracy

Memory performance (proportion correct) across each of the four experimental conditions are presented in Table 2.

**Table 2.** Mean proportion correct (and standard deviations) and mean proportion of joint retrieval (and standard deviations) for the data and independent model at no-delay (i.e., encoded immediately prior to retrieval delay) and delay (i.e., encoded 24 hours prior to retrieval) for closed- and open-loops.

| Delay | Loop | Proportion<br>Correct | Proportion of Joint Retrieval |  |
| --- | --- | --- | --- | --- |
|  |  |  | Data | Independent<br>Model |
| No-Delay | Open | 0.62 (0.19) | 0.61 (0.14) | 0.59 (0.12) |
|  | Closed | 0.68 (0.21) | 0.69 (0.14) | 0.65 (0.14) |
| Delay | Open | 0.50 (0.19) | 0.56 (0.13) | 0.55 (0.11) |
|  | Closed | 0.61 (0.21) | 0.67 (0.12) | 0.61 (0.13) |

A 2×2 (Loop x Delay) ANOVA revealed a significant effect of Loop, *F*(1, 49) = 37.72, *p* < .001, *η_p_^2^* = .44, and Delay, *F*(1, 49) = 45.74, *p* < .001, *η_p_^2^* = .48, with greater memory for closed-loops, relative to open-loops, and for pairs retrieved immediately after encoding, compared to those retrieved following a 24-hour delay. The ANOVA also revealed a significant interaction between these factors, *F*(1, 49) = 10.78, *p* = .002, *η_p_^2^* = .18, with a greater difference in memory between closed- and open-loops at delay relative to no-delay. This is consistent with finding reported in Joensen et al. (2020).

#### Retrieval dependency

The mean proportion of joint retrieval in the data and independent model, for closed- and open-loops at delay and no-delay, are presented in Table 2. Figure 2 shows mean retrieval dependency (i.e., data - independent model) for each condition. Evidence for dependency was seen for closed-loops at both delay, *t*(49) = 7.83, *p* < .001, *d* = 1.11, and no-delay, *t*(49) = 5.03, *p* < .001, *d* = .71. No evidence for dependency was observed for open-loops at delay, *t*(49) = 0.63, *p* = .533, *d* = .10, however we did find evidence for dependency for open-loops at no-delay, *t*(49) = 2.19, *p* = .033, *d* = .31 (but this effect did not survive corrections for multiple comparisons; adjusted alpha for four comparisons = .013), see Figure 2, though we note the smaller effect size estimate (0.31) relative to the two closed- loop conditions (1.11 and 0.71)

**Figure 2.**
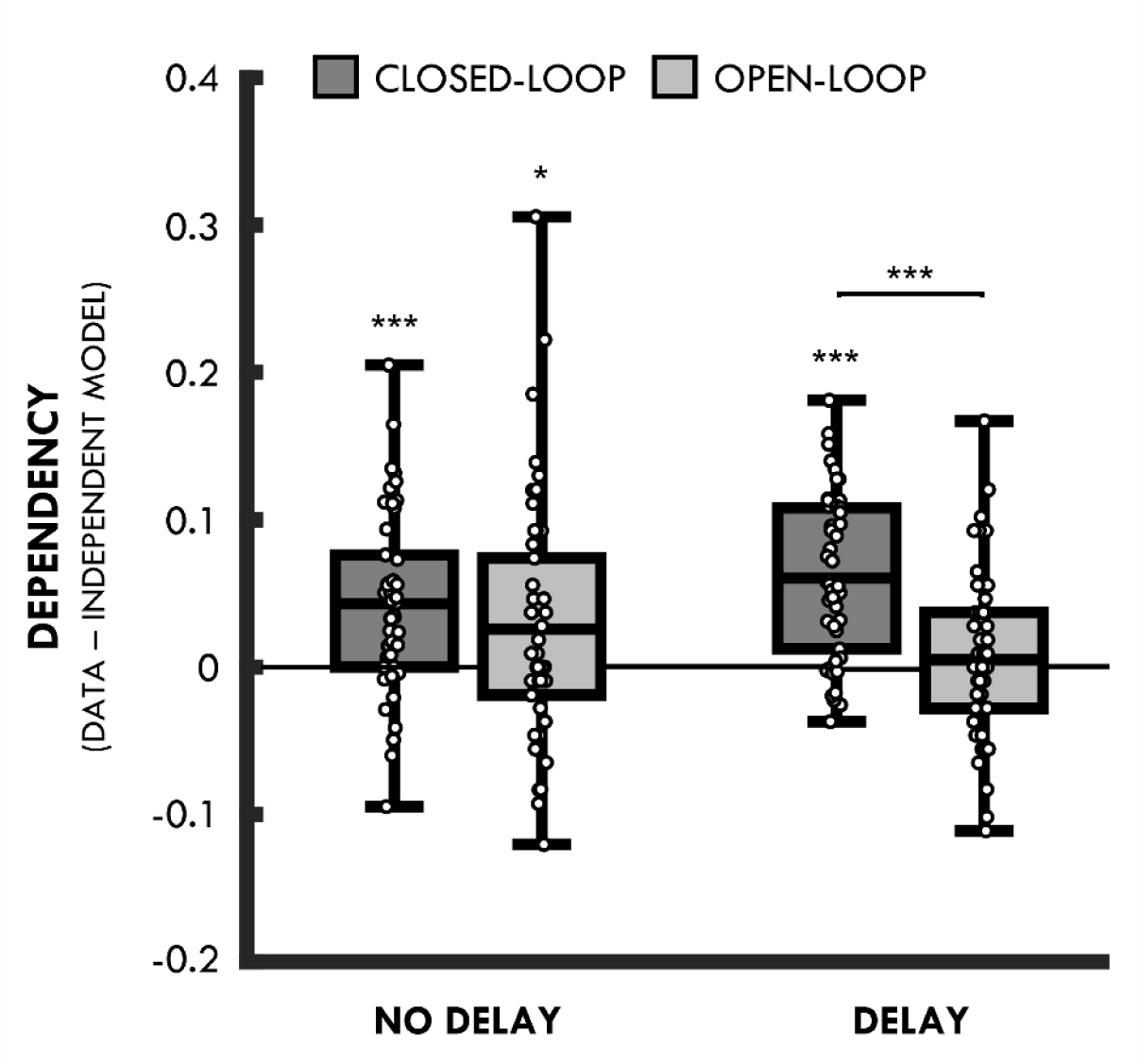
Dependency for closed- and open-loops at no-delay (i.e., encoded immediately prior to retrieval) and delay (i.e., encoded 24 hours prior to retrieval). Lines in boxes represent mean dependency in each condition. Bottom and top edges of boxes indicate the 25th and 75th percentiles, respectively. Whiskers represent the minimum and maximum data points. *Ns* = Not significant, *** *p* < .001, * *p* < .05.

A 2×2 (Loop x Delay) ANOVA revealed a significant main effect of Loop, *F*(1, 49) = 12.33, *p* < .001, *η_p_^2^* = .20, where dependency was greater for closed-compared to open-loops. However, this analysis also revealed a significant interaction, *F*(1, 49) = 5.21, *p* = .027, *ηp^2^* = .10, driven by a greater difference between closed- and open-loops at delay, *t*(49) = 5.03, *p* < .001, *d* = .71, relative to no-delay, *t*(49) = 1.13, *p* = .265 *d* = 0.16. Thus, we replicate previous findings showing greater dependency in the closed-loop delay than open-loop delay condition, but unexpectedly did not see the same pattern in the no-delay condition. Despite this, a direct comparison between the delay and no-delay open-loop condition was not significant, *t*(49) = 1.49, *p* = .142, *d* = 0.21, nor closed-loop condition, *t*(49) = 1.91, *p* = .062, *d* = 0.27. There was no main effect of Delay in this analysis, *F*(1, 49) = 0.04, *p* = .842, *η_p_^2^* < .01.

The finding of dependency in the open-loop delay condition was unexpected given previous studies have not found evidence for dependency for open-loops in an immediate test condition (Horner & Burgess, 2014; Horner et al., 2015; Joensen et al., 2020). One possibility is that the two encoding sessions in the present study (cf. Joensen et al. (2020) who had a single encoding session and two separate retrieval sessions) allowed participants to infer the associative nature of the overlapping pairs in the first encoding session (i.e., delay), leading to increases in dependency for open-loops encoded during the second encoding session (i.e., no-delay). However, it is important to note that dependency is seen for closed-loops at both delay and no-delay (i.e., it is the open-loop no-delay condition that appears to diverge from previous findings).

### fMRI

#### Neocortical Reinstatement

To look for differences in reinstatement as a function of Loop or Delay, we first identified regions where BOLD activity differed between the retrieval of different element types (i.e., people, locations, and objects), collapsed across the closed- vs open-loop and delay vs no-delay conditions. For example, to identify regions associated with the retrieval of people, retrieval trials where a person was either a cue or target were compared with trials where a person was neither the cue nor target (i.e., the non-target). The largest difference in BOLD activity for people was seen in the precuneus, for locations in the left parahippocampal gyrus, and for objects in the left middle temporal gyrus (see Table 3 & Figure 3A). Thus, as in Horner et al. (2015), people, locations and objects produced different levels of activation in specific neocortical regions. Unthresholded statistical maps of these effects are available at https://neurovault.org/collections/IFZTMWIU/.

**Figure 3.**
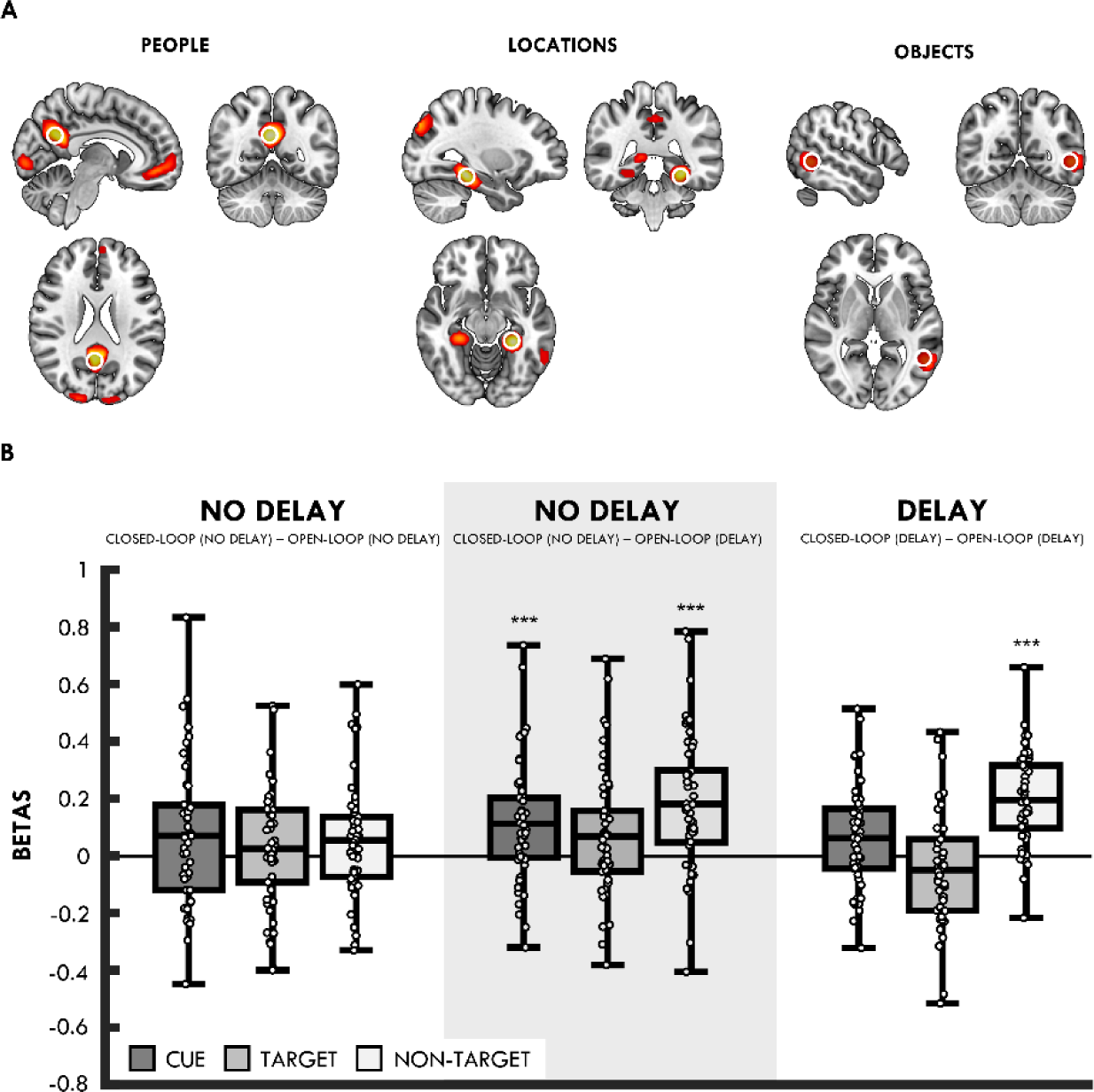
**(A)** Cortical regions showing activation differences for people, locations and objects. **(B)** Mean difference (collapsed across the precuneus, left parahippocampal gyrus and left middle temporal gyrus) in BOLD activity for closed- vs open-loops for cue (dark grey), target (light grey) and non-target (white) trials at no-delay, no delay (when using the open-loop condition at delay as baseline; highlighted with grey, transparent box), and delay. Lines in boxes represent mean difference in each condition. Bottom and top edges of boxes indicate the 25th and 75th percentiles, respectively. Whiskers represent the minimum and maximum data points. *** *p* < .001, corrected for multiple comparisons.

**Table 3.** Clusters and peaks showing element-specific whole brain activity for people, locations and objects (*p_FWE_* < .05; cluster size > 30. L = Left hemisphere; R = R hemisphere.

| Region | Voxels | MNI Coordinates |  |  | Z score |
| --- | --- | --- | --- | --- | --- |
|  |  | X | Y | Z |  |
| **People** |  |  |  |  |  |
| Precuneus Cortex | 450 | 3 | -57 | 27 | > 8 |
| L Lingual Gyrus | 1163 | -15 | -90 | -3 | > 8 |
| Frontal Medial Cortex | 368 | -3 | 48 | -9 | > 8 |
| L Middle Temporal Gyrus | 158 | -63 | -6 | -15 | > 8 |
| R Middle Temporal Gyrus | 56 | 63 | -3 | -18 | > 8 |
| Superior Frontal Gyrus | 61 | -6 | 57 | 30 | 7.80 |
| **Locations** |  |  |  |  |  |
| L Parahippocampal Gyrus | 270 | -27 | -36 | -15 | > 8 |
| L Lateral Occipital Cortex | 276 | -39 | -81 | 33 | > 8 |
| R Precuneus | 423 | 15 | -48 | 9 | > 8 |
| R Parahippocampal Gyrus | 159 | 27 | -30 | -18 | > 8 |
| R Lateral Occipital Cortex | 71 | 48 | -72 | 30 | 7.67 |
| R Cingulate Gyrus | 41 | -3 | -36 | 42 | 5.97 |
| L Middle Temporal Gyrus | 48 | -57 | -51 | -9 | 5.97 |
| L Superior Frontal Gyrus | 40 | -24 | 12 | 54 | 5.65 |
| **Objects** |  |  |  |  |  |
| L Middle Temporal Gyrus | 195 | -51 | -54 | 0 | 7.80 |
| L Frontal Pole | 30 | -48 | 39 | 12 | 5.63 |

BOLD responses from these regions were then extracted and assessed in terms of how they differed depending on whether the element associated with that region was either a cue, target, or non-target (e.g., a trial where an object was the cue and a location was the target would be a non-target trial for people). If closed-loops are reinstated holistically (regardless of delay), the region associated with the non-target (e.g., the precuneus for people) should show greater activity for closed-relative to open-loops (as non-target reinstatement is not expected to occur for open-loops).

We first tested whether we saw a non-target reinstatement effect, collapsed across the delay and no-delay condition and all three ROIs. This revealed significantly greater BOLD activity for closed-than open-loops in the non-target region, *t*(49) = 6.02, *p* < .001, *d* = .85. We also saw significantly greater BOLD response for closed-than open-loops in the cue region, *t*(49) = 2.92, *p* = .005, *d* = .41, but not the target region, *t*(49) = 0.61, *p* = .542, *d* = .09 (adjusted alpha for three comparisons = .017). We therefore provide evidence for non-target reinstatement as seen in previous studies (Horner et al., 2015; Grande et al., 2019).

We next tested whether the non-target reinstatement effect differed between the delay and no-delay conditions (Figure 3B). This revealed that BOLD activity for closed-relative to open-loops in the non-target region was significantly greater in the delay than no-delay condition, *t*(49) = 4.65, *p* < .001, *d* = .66. Interestingly, although we saw significantly greater BOLD activity for closed-than open-loops in the non-target region at delay, *t*(49) = 8.60, *p* < .001, *d* = 1.22 (adjusted alpha for two comparisons = .025), no difference was seen in the no-delay condition, *t*(49) = 1.89, *p* = .064, *d* = .27; though we note the numerical difference is in the same direction.

For completeness, we also tested for differences between closed- and open-loops in the cue and target region in the delay and no-delay condition. No differences between closed- and open-loops were seen in the cue region, delay: *t*(49) = 2.42, *p* = .020, *d* = .34; no-delay: *t*(49) = 2.01, *p* = .050, *d* = .29, or the target region, delay: *t*(49) = 1.76 *p* = .084, *d* = .25; no-delay: *t*(49) = 0.84, *p* = .405, *d* = .12 (adjusted alpha for four comparisons = .013).

Previous studies (Horner et al., 2015; Grande et al., 2019) observed a non-target reinstatement effect for closed-loops retrieved immediately after encoding, and as such the failure to replicate this effect was unexpected. However, the absence of a difference between closed- and open-loops at no-delay is perhaps consistent with the lack of a significant behavioural difference in dependency between closed- and open-loops at no-delay. Given that we observed evidence of dependency for open-loops at no-delay, it is possible that this open-loop condition is obscuring a possible non-target reinstatement effect in the closed-loop condition. To assess this possibility, we first directly compared the BOLD response in the delay and no-delay condition for the open-loop and closed-loop condition separately. This revealed significantly greater BOLD activity in the open-loop no--delay than delay condition in the non-target region, *t*(49) = 4.55, *p* < .001, *d* = .64 (adjusted alpha for six comparisons = .008), consistent with the possibility that participants are reinstating non-target elements more often in the open-loop no delay than delay condition. Apart from significantly greater BOLD activity in the closed-loop no-delay than delay condition in the target region, *t*(49) = 3.24, *p* = .002, *d* = .46), no further comparisons were significant, *t*s < 1.65, *p*s > .0105.

Given evidence that the BOLD activity in the open-loop no-delay condition differed from the open-loop delay condition in the non-target region, alongside the behavioural evidence for dependency in the open-loop no-delay condition, it is possible this condition is not serving as an appropriate baseline for comparison with the closed-loop no-delay condition. Given the lack of dependency in the open-loop delay condition, we reasoned this might be a more appropriate baseline for assessing non-target reinstatement in the no-delay condition.

The comparisons between the closed-loop no-delay and open-loop delay conditions revealed that BOLD activity was significantly greater for non-target closed-loops, relative to open-loops, *t*(49) = 5.33, *p* < .001, *d* = .76 (adjusted alpha for three comparisons = .017) (see Figure 3B). There was also a significant difference for cue, *t*(49) = 3.74, *p* < .001, *d* = .53, but not target trials when corrected for multiple comparisons, *t*(49) = 2.19, *p* = .034, *d* = .31. Thus, when we use a baseline in which we did not find evidence of behavioural dependency for open-loops (i.e., the open-loop delay condition), we see evidence for non-target reinstatement for closed-loops in both the delay and no-delay condition.

We therefore provide evidence for non-target reinstatement in the closed-loop delay and no-delay condition, although the effect at no-delay is only seen in a post-hoc analysis using the open-loop delay condition as a baseline measure. Critically, this post-hoc analysis does not affect the clear finding of non-target reinstatement following a 24-hour delay (including a night of sleep).

#### Non-target reinstatement and correlated activity

The above analyses provided evidence for non-target neocortical reinstatement in the closed-loop relative to open-loop condition after a 24-hour delay. This is consistent with behavioural evidence showing that closed-loops retain their coherence, as measured by retrieval dependency, following a delay (Joensen et al., 2020).

Previous work showed that the strength of non-target reinstatement correlated with the hippocampal closed- vs open-loop contrast across participants (Horner et al., 2015). This is consistent with the proposal that hippocampal pattern completion drives the reinstatement of all associated elements in the neocortex. To assess whether the role of hippocampal pattern completion differs over time, we correlated the non-target reinstatement effect (closed-vs open-loops) with the difference in BOLD activity between the closed- and open-loop condition across participants (as in Horner et al., 2015), separately for the no-delay and delay condition. Note, for this analysis we used the open--loop no-delay condition as the baseline measure for the closed-loop no-delay condition, despite this contrast not providing clear evidence for non-target reinstatement in the primary analysis.

At both delay and no-delay, multiple brain regions that have previously been associated with recollection, including the hippocampus (see Kim, 2010; Rugg & Vilberg, 2013; Spaniol et al., 2009 for reviews), correlated with the difference in non-target reinstatement between closed- and open-loops on a whole-brain level (see Table 4). Within the bilateral hippocampal mask (*p_FWE/_*_SVC_ < .05), we observed differences in BOLD activity for non -target trials that correlated with the closed-vs open-loop contrast at both delay and no-delay (see Table 5). We therefore provide evidence that non-target reinstatement correlates the hippocampal closed-vs open-loop constrast in both the no-delay and delay condition. Unthresholded statistical maps of these effects are available at https://neurovault.org/collections/IFZTMWIU/.

**Table 4.**
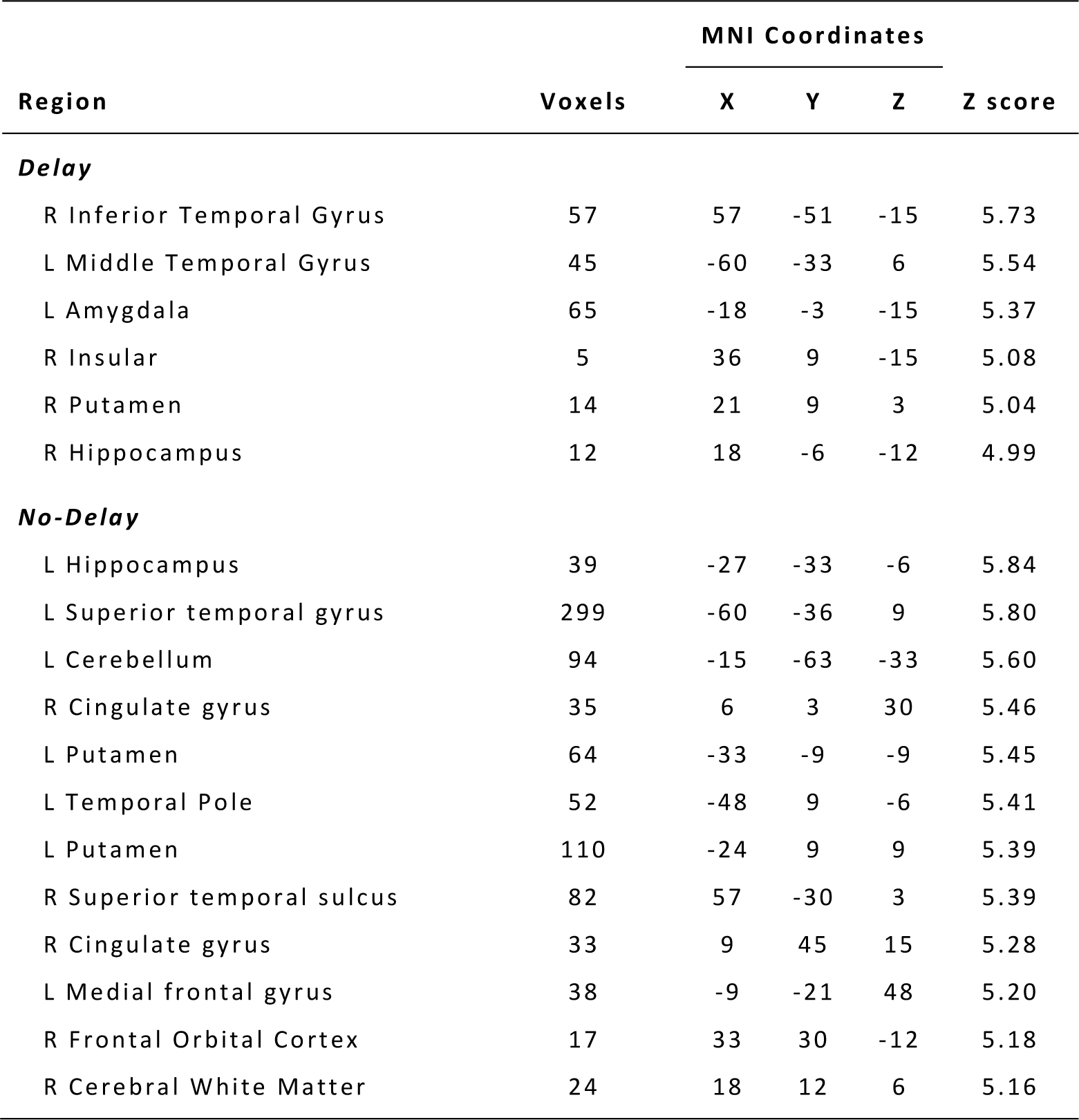
Clusters and peaks showing whole-brain regions, and clusters within a bilateral hippocampal mask, correlating with differences in cortical activity for non-target trials between closed- and open-loops at delay and no-delay (*p_FWE_* < .05, cluster size > 5). L = Left hemisphere; R = Right hemisphere.

**Table 5.** Clusters and peaks showing regions within the bilateral hippocampal mask correlating with differences in cortical activity for non-target trials between closed- and open-loops at delay and no-­delay (*p_FWE/SVC_* < .05, cluster defining threshold at the whole -brain level p < .001, cluster extent threshold > 5 voxels). L = Left hemisphere; R = Right hemisphere.

| Region | Voxels | MNI Coordinates |  |  | Z score |
| --- | --- | --- | --- | --- | --- |
|  |  | X | Y | Z |  |
| Delay |  |  |  |  |  |
| R Hippocampus | 2 | 18 | -6 | -12 | 4.99 |
| L Hippocampus | 1 | -18 | -6 | -12 | 4.79 |
| No-Delay |  |  |  |  |  |
| L Hippocampus | 26 | -27 | -33 | -6 | 5.84 |
| L Hippocampus | 1 | -33 | -12 | -12 | 4.80 |
| L Hippocampus | 4 | -21 | -9 | -12 | 4.74 |

We also performed a second analysis to assess the relationship between non-target reinstatement and the hippocampal closed- vs open-loop contrast. We first identified regions within the hippocampal mask that showed greater BOLD activity for closed-relative to open-loops (collapsed across delay and no-delay and all retrieval trials). This revealed a region in the left hippocampus (peak: −21, −36, 3) at an uncorrected threshold (*p* < .001) (Unthresholded statistical map of this effect is available at

https://neurovault.org/collections/IFZTMWIU/). We then extracted the difference between closed- and open-loops in this functionally defined region for each participant and correlated this difference with the difference in neocortical non-target activity between closed- and open-loops. Consistent with the whole-brain and SVC analyses reported above, we saw that activity from the closed- vs open-loop contrast in the hippocampus correlated with participants’ non-target reinstatement at both no-delay, *r* = 0.68, *p* < .001 (and when using the open-loop delay condition as baseline for the closed- loop condition at no-delay, *r* = 0.66, *p* < .001) and delay, *r* = 0.43, *p* = .002.

As a next step, to assess whether the relationship between the hippocampal closed- vs open-loop contrast and non-target reinstatement differed at no-delay and delay, we again extracted the difference between closed- and open-loops (collapsed across all retrieval trials) in the functionally defined hippocampal region identified above for each participant, as well as the mean difference in neocortical non-target activity between closed- and open-loops. We then used a generalised linear mixed-effects (GLME) model to estimate the strength of neocortical reinstatement (i.e., differences in neocortical non-target activity between closed- and open-loops), incorporating fixed effects of Delay (i.e., no-delay vs delay), the hippocampal closed- vs open-loop contrast (i.e., difference in hippocampal activity between closed- and open-loops), and their interaction, along with random effects of participant and Delay for each participant. Note that the fixed-effects specify predictors of interest, while the random effects account for statistical dependency between within-participant observations.

Consistent with the correlation analyses, the GLME model revealed significant effects of the hippocampal closed- vs open-loop contrast on the strength of non-target reinstatement at both no-delay, *t*(96) = 6.49, *p* < .001, and delay, *t*(96) = 3.43, *p* < .001 (adjusted alpha for six comparisons = .008). We also saw that the slope of this effect of the hippocampal contrast on non-target reinstatement did not differ significantly between delay and no-delay condition, *t*(96) = 1.62, *p* = .11. As such, the hippocampal closed- vs open-loop contrast is predictive of non-target reinstatement in both the no-delay and delay conditions, and we see no evidence for a difference across delay.

Critically, the GLME model revealed that the intercepts differed significantly across delay, *t*(96) = 4.78, *p* < .001. While the intercept at no-delay did not differ from zero, *t*(96) = 1.32, *p* = .189, the intercept for the delay condition was significantly greater than zero, *t*(96) = 6.82, *p* < .001 (see Figure 4). This is critical as it demonstrates that, although the hippocampal closed- vs open-loop contrast correlated with non-target reinstatement at both time points, the reinstatement effect can occur without evidence for a corresponding difference between closed- vs open-loops in the hippocampus at delay. Given this hippocampal contrast is likely a marker of hippocampal pattern completion (Horner et al., 2015), this suggests that non-target reinstatement can be driven, in an additive manner, by both hippocampal pattern completion and a non-hippocampal process following a 24-hr delay between encoding and retrieval.

**Figure 4.**
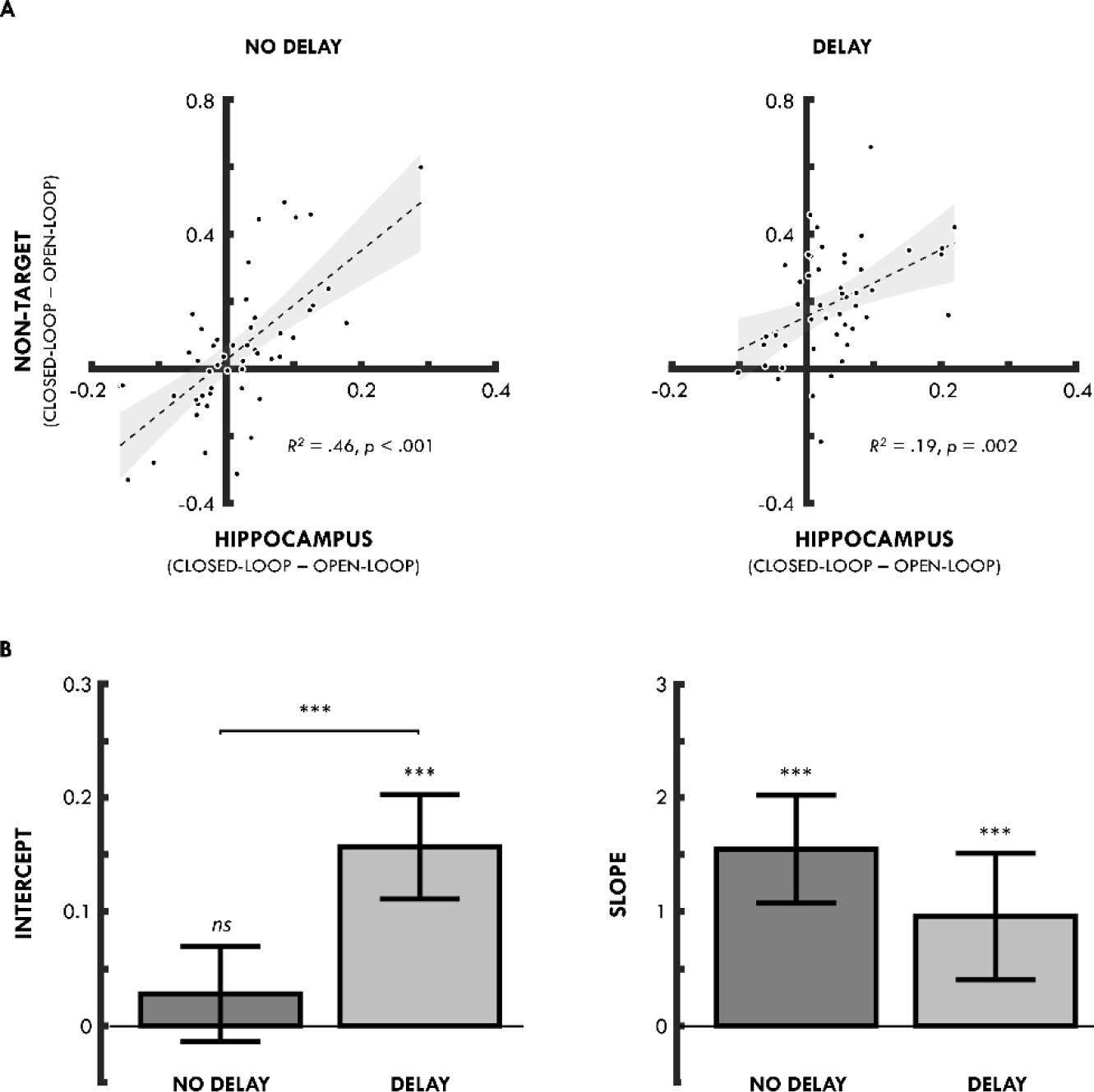
**(A)** Correlation between the difference in hippocampal activity during the retrieval of closed- vs open-loops (hippocampal ROI defined as a 9-mm radius sphere centred on the peak coordinate from the closed- vs open-loop contrast peak: −21, −36, 3) and the mean difference (collapsed across precuneus, left parahippocampal gyrus and left middle temporal gyrus for people, locations and objects, respectively) in BOLD activity during non-target retrieval for closed- vs open-loops, separately for events retrieved after no--delay (left) and delay (right). **(B)** Mean GLME model estimates (and 95% confidence intervals) for intercept (left) and slope (right) for the no-delay (dark grey) and delay (light grey) conditions. *** p < .001, *ns* non-significant.

To assess the robustness of these effects, we re-ran the GLME analysis while removing a randomly selected subset of participants at each analysis point from *n*-1 to *n*-20. For each of these analyses, we iteratively repeated the analysis 50 times and removed a randomly selected subset of participants for each iteration. The robustness analysis showed that the significant intercept differences from zero in the delay condition was present in every iteration, as well as the significant differences in the intercept between the no-delay and delay condition being present in every iteration. The non-significant differences in the slope between the no-delay and delay condition was more variable, decreasing below 90% consistency with the main analysis at n-9, with a mean consistency of 83% between *n*-9 and *n*-20 and a minimum consistency of 70% at *n*-20. Thus, the results of our novel GLME approach to assessing the relationship between hippocampal pattern completion and non-target neocortical reinstatement are robust to random removal of data points to at least *n*-9 (i.e., a total *N* of ∼40).

## Discussion

We show that events, composed of three overlapping pairs (i.e., closed-loops), continue to be reinstated holistically following a 24-hr period of forgetting and consolidation. Although memory performance decreased over time (indicating that forgetting had occurred), we continued to observe BOLD activity related to the reinstatement of elements incidental to retrieval (i.e., non-targets). For example, when participants were cued with a location (e.g., *kitchen*) and asked to retrieve the overlapping object (e.g., *hammer*), retrieval also led to activity in cortical regions associated with the overlapping person (e.g., *Barack Obama*), despite no explicit demand to retrieve this element (i.e., the element was neither the cue nor the retrieval target). Importantly, the level of this non-target reinstatement effect at delay was equivalent to that seen for closed-loops retrieved immediately after encoding (when using the open-loop condition at delay as baseline). This is consistent with behavioural evidence for ‘holistic forgetting’ where events retain their coherence over periods of forgetting (Joensen et al., 2020).

The reinstatement of non-target elements is thought to be a direct consequence of pattern completion in the hippocampus (Horner & Doeller, 2017) whereby a partial input can result in the retrieval of the complete memory. We show that the strength of the non-target reinstatement effect correlates with a signature of hippocampal pattern completion (hippocampal closed- vs open-loop contrast) before and after a 24-hr delay. However, although the slope of this relationship did not statistically differ across delay, the intercept significantly increased and was statistically greater than zero at delay. Although this novel analysis and finding appears to be robust to the removal of random participants (at least to N-9 participants), it is nonetheless important that they should be replicated in an independent data set in the future. Our results suggest that hippocampal pattern completion still contributes to the strength of this effect after delay, however some non-target reinstatement can occur without a corresponding signature of hippocampal pattern completion following a 24-hr delay. This latter finding points to an emergent non-hippocampal contribution to holistic reinstatement.

It has long been proposed that the role of the hippocampus in long-term memory retrieval changes over time (McClelland et al., 1995; Squire & Alvarez, 1995). This proposal is motivated by observations that although patients with hippocampal damage present with amnesia (Scoville & Miller, 1957; Vargha-Khadem et al., 1997), they often show a temporal gradient to their deficit, with memory for remote events being relatively spared compared to more recent ones (Schnider et al., 1995; but see also Sanders & Warrington, 1971).

Standard theories of consolidation have argued that the sparse activity patterns and high neural density of the hippocampus are thought to provide the ideal mechanisms for the initial encoding and retrieval of event-based representations (McClelland et al., 1995; Norman & O’Reilly, 2003; Treves & Rolls, 1994). However, over periods of consolidation, gradual adjustments to cortical connections are thought to allow for the formation of neocortical representations that can be retrieved independent of the hippocampus (McClelland et al., 1995). Although, standard theories of consolidation have noted that this process can be relatively protracted occurring over days, weeks, or months, there is evidence that differences in hippocampal activity at retrieval already emerge after a one- day interval (Takashima et al., 2006; Takashima et al., 2009).

Consistent with these findings, we found a difference in the intercept between the no-delay and delay conditions when estimating the relationship between neocortical reinstatement and hippocampal pattern completion. While the intercept did not differ from zero in the no-delay condition, it was significantly greater than zero in the delay condition (and the intercepts significantly differed between conditions). The non-zero intercept suggests that neocortical reinstatement can occur without evidence for a corresponding signature of hippocampal pattern completion following a 24-hr period of consolidation. Yet, our results also show that while the neocortical reinstatement of non-target elements can be achieved without a corresponding increment in hippocampal pattern completion, the hippocampus still contributes to the strength of this reinstatement effect. As such, while consolidation may allow for the strengthening of related neocortical representations that can be holistically reinstated independently of hippocampal pattern completion - consistent with standard theories of consolidation - pattern completion continues to mediate the strength to which events are holistically reinstated after a 24-hr delay.

Critically, these two contributions to holistic neocortical reinstatement appear to be additive in nature; the total amount of reinstatement is the sum of the linear relationship between hippocampal pattern completion and reinstatement and the additional non- hippocampal contribution (at least after 24-hours). This suggests that the relationship is not compensatory in nature - it isn’t the case that as one contribution increases the other decreases.

One critical question is whether this pattern is consistent across further delays. It is possible that the hippocampal contribution decreases over longer timescales, which would be reflected in a decrease in the slope of the estimated relationship over time. Such a result would support standard consolidation theories; however, it would point to a more additive relationship during the consolidation process, with hippocampal pattern completion continuing to contribute to the reinstatement of past events even in the presence of a non-hippocampal contribution to retrieval. Indeed, our approach potentially allows for the tracking of independent time courses, where the contribution of hippocampal pattern completion might decrease over time at a different rate relative to the emergence of the non-hippocampal process, and that these different time courses might be driven by different mechanisms. For example, the increase in the non- hippocampal process might be driven by active systems consolidation processes during sleep (Diekelmann & Born, 2010), whereas the decreased contribution of hippocampal pattern completion might be a function of forgetting via decay mechanisms (Hardt et al., 2013; Sadeh et al., 2013). Tracking the time course of changes to the slope and intercept of the hippocampal pattern completion-neocortical reinstatement relationship over different delays (from one day to several weeks) could assess this possibility.

Alternative accounts of consolidation, such as the multiple trace theory (Nadel & Moscovitch, 1997) and trace-transformation theory (Sekeres et al., 2018; Winocur & Moscovitch, 2011) make a distinction between the maintenance and retrieval of fine-grained, or context-rich, information and more gist-like information surrounding a memory. These distinct representations have been proposed to be represented independently, with the former being more dependent on the hippocampus and the latter on the neocortex. Support for this distinction has come from evidence showing that peripheral details are forgotten more rapidly than gist-like information from an event (Sekeres et al., 2016) and evidence showing that memory for context-rich information of an event (but not gist-like information) tends to be impaired following hippocampal damage (St-Laurent et al., 2009).

Our findings may represent an intermediate between standard theories of consolidation and these alternative accounts, where the direct involvement of hippocampal pattern completion in recollection endures, insofar as hippocampal pattern completion still contributes to the strength or precision of reinstatement, despite the emergence of a non-hippocampal contribution to retrieval. Harlow and colleagues (Harlow & Donaldson, 2013: Harlow & Yonelinas, 2016) have argued that recollection may be characterised by two potentially separable memory components: ‘accessibility’ (i.e., the probability of successful retrieval) and ‘precision’ (i.e., the precision/fidelity of the successfully retrieved information). As such, it is possible that the strength (or precision) of reinstatement is reflective of a more continuous retrieval process, such that even in instances where an entire event is successfully reinstated in the neocortex, the fidelity of reinstatement may differ across events, and this may depend on the involvement of the hippocampus (e.g., Korkki et al., 2021; Nilakantan et al., 2018, but see also Richter et al., 2016). Whereas some initial (less precise) neocortical reinstatement could occur without hippocampal pattern completion following a delay, further (more precise, detailed) reinstatement could still require pattern completion. This would predict that when retrieving context-rich information the hippocampal-neocortical relationship should be maintained over delay, however, when retrieving more gist-like information, the relationship should decrease over delay. Assessing memory for the perceptual versus semantic features of multiple event elements across delay could address this possibility.

In sum, we provide evidence for holistic reinstatement following a period of forgetting and consolidation. This is consistent with previous behavioural evidence suggesting that event-based memories undergo a holistic form of forgetting where events that are retained continue to be retrieved in their entirety and events that are forgotten are forgotten in their entirety. Critically, we found evidence that holistic reinstatement in the neocortex is driven by both hippocampal pattern completion and a non-hippocampal process after 24-hours. Further, our novel experimental approach allows us to independently assess both a hippocampal and non-hippocampal contribution to reinstatement, allowing future research to track the time-courses of these potentially independent contributions. Finally, our results show that holistic retrieval is driven by both a non-hippocampal process *and* hippocampal pattern completion after 24-hours, suggesting that the hippocampus and neocortex work in an additive, rather than compensatory, manner to support episodic memory retrieval.

## Data availability

Second level data and analyses are available on the Open Science Framework (https://osf.io/hk79v/).

